# Cooperativity between Cas9 and hyperactive AID establishes broad and diversifying mutational footprints in base editors

**DOI:** 10.1101/2022.12.03.518995

**Authors:** Kiara N. Berríos, Aleksia Barka, Jasleen Gill, Juan C. Serrano, Peter F. Bailer, Jared B. Parker, Niklaus H. Evitt, Kiran S. Gajula, Junwei Shi, Rahul M. Kohli

**Affiliations:** Graduate Group in Biochemistry and Molecular Biophysics, Perelman School of Medicine, University of Pennsylvania, Philadelphia, Pennsylvania, 19104, USA; Department of Medicine, Perelman School of Medicine, University of Pennsylvania, Philadelphia, Pennsylvania, 19104, USA; Graduate Group in Cell and Molecular Biology, Perelman School of Medicine, University of Pennsylvania, Philadelphia, Pennsylvania, 19104, USA; Department of Cancer Biology, Perelman School of Medicine, University of Pennsylvania, Philadelphia, Pennsylvania, 19104, USA

## Abstract

The partnership of DNA deaminase enzymes with CRISPR-Cas nucleases is now a well-established method to enable targeted genomic base editing. However, an understanding of how Cas9 and DNA deaminases collaborate to shape base editor (BE) outcomes has been lacking. Here, we support a novel mechanistic model of base editing by deriving a range of hyperactive activation-induced deaminase (AID) base editors (hBEs) and exploiting their characteristic diversifying activity. Our model involves multiple layers of previously underappreciated cooperativity in BE steps including: (1) Cas9 binding can potentially expose *both* DNA strands for ‘capture’ by the deaminase, a feature that is enhanced by guide RNA mismatches; (2) after strand capture, the intrinsic activity of the DNA deaminase can tune window size and base editing efficiency; (3) Cas9 defines the boundaries of editing on each strand, with deamination blocked by Cas9 binding to either the PAM or the protospacer; and (4) non-canonical edits on the guide RNA bound strand can be further elicited by changing which strand is nicked by Cas9. Leveraging insights from our mechanistic model, we create novel hBEs that can remarkably generate simultaneous C>T and G>A transitions over >65 bp with significant potential for targeted gene diversification.

## INTRODUCTION

Cytosine base editors (BEs) combine the powerful DNA deamination activity of AID/APOBEC family enzymes with CRISPR-Cas targeting in order to introduce point mutations at a specified locus (1). In the first generation of base editors (BE1), rat APOBEC1 (rA1) was fused to a catalytically dead *S. pyogenes* Cas9 (dCas9) and targeted using a single-guide RNA (sgRNA) (2). Upon dCas9-sgRNA binding to the target strand (TS), R-loop formation exposes the non-target strand (NTS) resulting in single-stranded DNA (ssDNA) that is then ‘captured’ by the DNA deaminase. The NTS can then be deaminated in this exposed ‘editing window’, typically 12-16 base pairs (bp) away from the protospacer adjacent motif (PAM), and the resulting C/G>T/A mutations are installed after replication over the lesion.

Since BE1, several strategies have been employed to increase precise editing or alternatively to generate diversifying mutations at the target site (2–9). Major improvements to BEs that enhance efficiencies include: (1) the incorporation of UGI – a small phage-derived inhibitor of uracil DNA glycosylase – to suppress base excision repair in BE2 editors, (2) the replacement of dCas9 with a TS-nickase (Cas9-D10A, nCas9) to promote replication over NTS mutations in BE3 editors, and (3) mammalian codon optimization along with the addition of a second UGI in BE4max editors (2, 10, 11). Novel BEs with diverse features have also been developed by using deaminases beyond rA1, including human AID or APOBEC3 enzymes, as well as engineered TadA variants, allowing for cytosine base editors (CBEs) or adenine base editors (ABEs) (4, 12-15). Additionally, Cas9 engineering has been employed to alter outcomes, with PAM-modified or circularly permuted Cas9 variants increasing the targeting scope of base editors (16, 17).

Given the transformative potential of BEs, intense efforts have focused on the development of improved tools. While these remarkable advances have now yielded a broad and powerful toolbox for genome engineering, fundamental mechanistic questions remain unanswered. In particular, the critical question of how Cas9 and DNA deaminases coordinate to promote BE outcomes has yet to be fully explored. The current model for BE-mediated mutagenesis focuses on mutation of the NTS and involves multiple steps: Cas9 binding and unwinding of the DNA duplex, AID/APOBEC capture of the ssDNA bubble and enzymatic deamination, along with TS nicking and replication over the uracil lesion (Figure 1A). When considering this stepwise mechanism, illuminating the interplay of Cas9 engagement and DNA deaminase action in each step could facilitate the continued optimization and expansion of BEs and support the development of the next generation of genome engineering tools.

**Figure 1.**
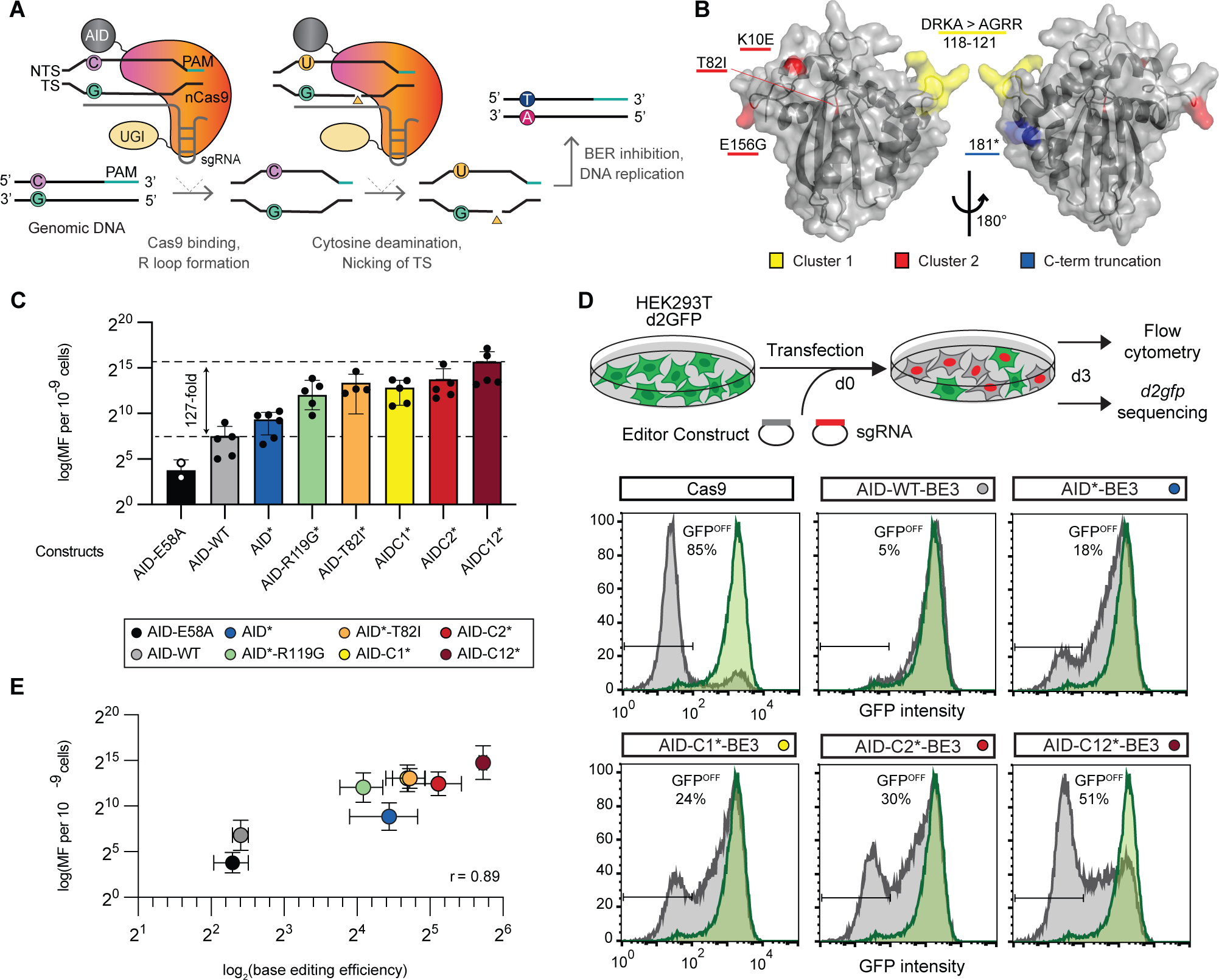
Hyperactive AID deaminases increase base editing activity. **(A)** In base editing, sgRNA-bound nCas9 binds the target strand (TS) forming an R loop exposing ssDNA on the non-target strand (NTS). The DNA deaminase enzyme then converts C>U bases in the exposed ssDNA. TS nicking by nCas9, inhibition of base excision repair (BER) pathway, and replication over the uracil locks in C/T to G/A mutations in genomic DNA. **(B)** Schematic of the AID structure(51) (PDB 5w0z) showing clusters of hyperactivating mutations tested in this study. **(C)** Mutation frequency, as measured by the frequency of acquired rifampin resistance (Rif^R^ mutation frequency, MF) upon expression of AID variants in *E. coli.* Individual data points (n = 2-6) are shown with mean and standard deviation noted, with individual values provided in Table S1. AID-E58A, catalytically inactive control. **(D)** Editing was evaluated in a HEK293T cell line containing *d2gfp*. The presence of *d2gfp*-targeting sgRNA can introduce indels or mutations to generate GFP^off^ cells, which can be tracked by flow cytometry. Shown are representative flow cytometry histograms from transfection of select constructs. Individual data points (n = 3) values are given in Table S1. **(E)** Correlation plot of intrinsic DNA deaminase activity as measured by Rif^R^ in (**C**) vs. BE activity with average and standard deviation from n≥3 replicates shown.

As noted, the existing toolbox for BEs has utilized diverse members of the AID/APOBEC family, each with distinctive sequence preferences, intrinsic deamination rates, cellular localization, and modes of action. As these features are retained when DNA deaminases are incorporated into BEs, it has been challenging to decipher the impact of *single* attributes on the BE mechanism. By keeping the BE scaffold the same and manipulating specific attributes, the underlying reaction mechanisms could potentially be better elucidated. In this regard, AID provides several distinctive features that make it an ideal candidate for systematic variation. AID naturally functions to drive somatic hypermutation (SHM) in B-cells where it generates clustered mutations on the immunoglobulin (Ig) locus by acting on either strand in a processive fashion (18, 19). In addition to being processive, AID activity can be tuned as its deamination activity has been highly constrained (20), as evidenced by the isolation of various hyperactivating mutations that shift the balance towards increased genomic instability in B-cells (21, 22). Similar to the opportunity offered by systematically altering AID, perturbing Cas9 cleavage activity can offer a means to alter downstream replication and repair, and analyzing BE off-targets can provide a means to assess the impact of Cas9 engagement. Thus, AID-focused BEs provide an ideal model system for understanding the role of each step in the BE mechanism.

In this study, we aimed to create a complete mechanistic model for how collaboration between Cas9 and AID establish BE outcomes. To this end, we derive a range of hyperactive AID base editors (hBEs), and reveal that BE activity and editing windows can be directly tuned by altering the intrinsic activity of the associated DNA deaminase. By further perturbing Cas9 engagement and downstream nicking and repair, hBEs provide unexpected new insights, revealing that both NTS and TS can be captured by the enzyme for subsequent deamination. Both mismatched sgRNAs and Cas9 nicking variants can increase unexpected editing at the TS, resulting in a striking increase in G>A mutations neighboring C>T changes. Beyond advancing a complete model for the impact of Cas9 binding, deaminase strand capture, deamination, and DNA repair in BEs, our work yields a simplified diversifying BE with a distinctive and novel capability of efficiently generating simultaneous C>T and G>A transitions over a >65 bp window. Our systematic approach shows how BE mutational landscapes are cooperatively established by Cas9 and AID and how modifying their activities can uncover new gene editing outcomes for diverse biotechnological applications.

## MATERIALS AND METHODS

### Design and cloning of hyperactive DNA deaminase constructs

For bacterial studies with AID enzymes, the pET41 expression plasmids containing AID-WT, AID-E58A, or AID* and an N-terminal maltose binding protein tag (MBP), which have been previously described (23), were used. For hyperactive AID constructs, AID inserts with C1* (D118A, R119G, K120R, A121R), C2* (K10E, T82I, E156G) and C12* (D118A, R119G, K120R, A121R, K10E, T82I, E156G) mutations were obtained as gBlocks (IDT) and cloned into the AID* containing parent construct. The R119G and T82I point mutations were added to the parent AID* construct using the Q5 Site-Directed Mutagenesis Kit (NEB, Cat # E0554S).

For BE3 base editing constructs, the various AID constructs from pET41 constructs were cloned into the scaffold of pCMV_BE3 (Addgene #73021), which contains rat APOBEC1. To this end, the parent pCMV_BE3 plasmid was digested with NotI and XmaI to remove rat APOBEC1. The AID coding sequences were amplified from their respective pET41 constructs, adding flanking regions of overlap with the pCMV_BE3 plasmid backbone. The final plasmids were then constructed using Gibson Assembly Master Mix (NEB, Cat # E2611S), merging the amplified gene fragments with the NotI/XmaI digested parent vector.

For the AID-C12*-nCas9-BE4max construct, AID’-BE4max (Addgene #174696) was used as a scaffold. This parent plasmid contained two point mutations, one in the linker region between Cas9 and UGI and the second one in the first UGI (E11K) protomer. These mutations were corrected here by ligating the NotI/SacI digestion fragment from pCMV-BE4max (Addgene #112093) into the digested AID’-BE4max (Addgene #174696) via T4 DNA ligase (NEB, Cat # M0202S). After generation of this AID-C12*-nCas9-BE4max construct, the point mutations to synthesize AID-C12*-dCas9-BE4max and AID-C12*-n’Cas9-BE4max were generated using the Q5 Site-Directed Mutagenesis Kit (NEB, Cat #E0554S).

The AID-C12*-nCas9-BE4max construct was also used as scaffold to generate AID-C12*-XTEN80-nCas9-BE4max by replacing the original 32aa-long SGGS-XTEN-SGGS linker between AID-C12* and nCas9 with the 80aa-long XTEN80 linker present in the epigenome editor TETv4 (24). Cloning of this construct was performed by GenScript.

For our R-loop assays, we constructed a constitutive all-in-one dead SaCas9 (dSaCas9) system from construct (Addgene #164563) via Gibson Assembly Master Mix (NEB, Cat # E2611S) where both dSaCas9 and its targeting-gRNA (targeting Chr 9: 21036-21332) are independently translated. This construct was then used to generate an all-in-one nickase SaCas9 (SaCas9-D10A, nSaCas9).

### Design and cloning of sgRNA expression plasmids

The sgRNA expression plasmids were generated using oligonucleotide cassettes and the LRcherry2.1 plasmid (25). Oligos containing the spacer sequence and Esp3I sticky ends were annealed and phosphorylated using T4 Polynucleotide Kinase (NEB) and purified using the Oligo Clean and Concentrator kit (Zymo Research). To generate mismatched sgRNAs, the spacer sequence targeting the *d2GFP* locus was modified by inserting mutations distal to the PAM. LRcherry2.1 plasmid was incubated with Esp3I (Thermo Fisher Scientific) at 37 °C for 2 hours to remove a short filler sequence, and further agarose gel purified. The sgRNA cassettes were then ligated in place of the filler using T4 DNA ligase (NEB).

### Bacterial DNA deaminase rifampin mutagenesis assay

The previously reported rifampin mutagenesis assay (23, 26) was used to measure the mutation frequency of AID variants. Plasmids encoding the AID variants (Kanamycin resistant) were transformed into BL21(DE3) *E. coli* harboring a plasmid encoding uracil DNA glycosylase inhibitor (UGI) (chloramphenicol resistant) (23). Overnight cultures were grown in LB with kanamycin (30 ng/mL) and chloramphenicol (25 ng/mL) from single colonies and diluted to an OD_600_ of 0.2. Cells were then grown for 1 hour at 37 °C until they reached OD_600_ of ∼0.6 and deaminase expression was induced with 1mM isopropyl β-D-1-thiogalactopyranoside (IPTG). After 4 hours of additional growth, serial dilutions were plated on Luria Bertani (LB) agar plates containing rifampin (100 μg/mL) and plasmid-selective antibiotics (kanamycin and chloramphenicol). The mutation frequencies were then calculated by the ratio of rifampin-resistant (Rif^R^) colonies relative to the total colony forming units.

### Cell culture

Previously derived HEK293T-d2GFP reporter cells (26) were cultured in Dulbecco’s Modified Eagle Medium (DMEM) media (Mediatech, Cat # MT10-013-CV) supplemented with 10% calf serum (CS; Fisher Scientific, Cat # SH3007203) and 1% penicillin/streptomycin (Invitrogen, Cat # 15140122) at 37 °C with 5% CO2 (26).

### Base editing assay using d2GFP inactivation in HEK293T cells

The base editing assay was modelled on a previously published protocol (26). Briefly, HEK293T-d2GFP cells were seeded on 24-well plates and transfected at approximately 60% confluency. 660 ng of the hBE constructs and 330 ng of LRcherry2.1 sgRNA expression plasmids were transfected using 1.5 µL of Lipofectamine 2000 CD (Invitrogen, Cat # 12-566-014). Negative control samples include LRcherry2.1T plasmid lacking a protospacer (labeled as no sgRNA samples). The *d2gfp-*targeting sgRNA can support the introduction of indels in *d2gfp* with Cas9 or a Q158X nonsense mutation with base editing, along with other potential edits. Transfected cells were harvested at day 3 after transfection, ensuring single-cell suspension. Genomic DNA was collected from cells using the DNeasy Blood & Tissue Kit (Qiagen, Cat # 69506) and the *d2GFP* locus was amplified and then deep sequenced as described on the “DNA library preparation” section below. Representative experiments were repeated independently three times and the results were reproducible.

### R-loop assay

The R-loop assay used in this study was based of a previously published protocol (26). Briefly, HEK293T-d2GFP cells were seeded on 24-well plates and transfected at ∼60% confluency. 400 ng of hBE constructs, 200 ng of *EMX1*-targeting LRcherry2.1 sgRNA plasmid, and 400 ng of dSaCas9 or nSaCas9 expression plasmid were co-transfected using 1.5 µL of Lipofectamine 2000 CD (Invitrogen, Cat # 12-566-014) per well. Transfected cells were harvested at day 3 after transfection, ensuring single-cell suspension. Genomic DNA was collected from cells using the DNeasy Blood & Tissue Kit (Qiagen, Cat # 69506) and both the *EMX1, EMX1-OT1, EMX1-OT2* and dSaCas9-targeted locus (Chr 9: 21036-21332) were amplified and then deep sequenced as described on the “DNA library preparation” section below. Representative experiments were repeated independently three times and the results were reproducible.

### Cell viability and DNA damage analysis of base edited HEK293T cells

For viability and DNA damage analysis, the base editing assays were performed as described above, with minor modifications (26). In these experiments, negative control samples include cells transfected with a LRcherry2.1T plasmid lacking a protospacer (labeled as no sgRNA samples). Positive control samples include cells transfected with 500 ng previously described pLEXm-APOBEC3A (27) or incubated with 5mM hydroxyurea. As above, transfected cells were harvested at day 3 after transfection. For cell viability analysis, a portion of cells were characterized using the LIVE/DEAD™ Fixable Green Dead Cell Stain Kit (ThermoFisher, Cat # L23101) per the manufacturer’s protocol. Another portion of cells was fixed, permeabilized, and stained with an AlexaFluor® 488 Mouse anti-γH2AX (pS139) antibody (BD Pharmingen™, Cat # 560445) per the manufacturer’s protocol. A total of 50, 000 cell events were counted on a Guava Easycyte 10HT instrument (Millipore, https://www.emdmillipore.com/US/en/20130828_204410). The GFP^on^ population gating, denoting γH2AX+ cells, was performed using the nadir between the two peaks on a sample transfected with APOBEC3A, with consistent gates for all comparison experiments. Representative experiments were repeated independently three times, and the results were reproducible. Flow cytometry analysis was performed using FlowJo Software Version 10.7.1 (FloJo, LCC, https://www.flowjo.com).

### Base editing of various genomic loci

The base editing assay for diverse genomic loci was based of a previously published protocol (26). Briefly, the transfection protocol was performed as above, but with different sgRNAs targeting various loci instead of the *d2gfp*-targeting sgRNA. Transfected cells were harvested at day 3 after transfection, ensuring single-cell suspension. Genomic DNA was collected using the DNeasy Blood & Tissue Kit (Qiagen, Cat # 69506) and each locus was amplified and then deep sequenced as described on the “DNA library preparation” section below. Representative experiments were repeated independently three times and the results were reproducible.

### DNA library preparation

Genomic DNA was collected from cells using the DNeasy Blood & Tissue Kit (Qiagen, Cat # 69506). The target loci of interest were PCR-amplified from 100 ng genomic DNA using KAPA HiFi HotStart Uracil+ Ready Mix (Kapa Biosystems, Cat # KK2602) or Phusion High-Fidelity DNA Polymerase (New England Biolabs, NEB, Cat # M0531S). PCR products were purified via QIAquick PCR Purification Kit (Qiagen, Cat # 28106). Indexed DNA libraries were prepared using the NEBNext Ultra II DNA Library Prep Kit (NEB, Cat # E7645S) with the following specifications. After adapter ligation and 4 cycles of PCR enrichment, indexed amplicon concentration was quantified by Qubit dsDNA HS Assay Kit (ThermoFisher, Cat # Q32854). Indexed PCR amplicons were pooled together in an equimolar ratio for paired-end sequencing by MiSeq (Illumina) with the 300-cycle MiSeq Reagent Nano Kit v2 (Illumina, Cat # MS-103-1001) or 300-cycle MiSeq Reagent Micro Kit v2 (Illumina, Cat # MS-103-1002).

### Statistics

Statistics were performed using GraphPad Prism. Comparisons of > 2 groups was performed using one-way ANOVA adjusted for Tukey’s multiple comparisons. Information on statistical tests performed, exact values of n, and how significance was defined is available in the figure legends or provided as Supplementary Data. For cell culture experiments, n is defined as cell cultures treated with separately prepared transfection solutions in different days.

### Flow cytometry

For GFP editing analysis, HEK293T-d2GFP cells were harvested and 50, 000 total cell events were gated with FSC/SSC gate on a Guava Easycyte 10HT instrument (Millipore). A second gate for high mCherry transfection efficiency was applied. The GFP^off^ population gating was performed using the nadir between the two peaks, with consistent gates for all comparison experiments. Flow cytometry analysis was performed using FlowJo Software Version 10.7.1 (FloJo, LCC, https://www.flowjo.com).

### Next-Generation Sequencing Analysis

Raw reads were automatically demultiplexed by MiSeq Reporter. Demultiplexed read qualities were evaluated by FastQC v0.11.9 (http://www.bioinformatics.babraham.ac.uk/projects/fastqc/). Low-quality sequence (Phred quality score <28) and adapters were trimmed via Trim Galore v0.6.5 (http://www.bioinformatics.babraham.ac.uk/projects/trim_galore/) prior to analysis with CRISPResso2 (28) Sequencing yielded ∼1, 000-50, 000 aligned reads per sample.

## RESULTS

### Modulating intrinsic AID activity tunes BE activity

To first generate a potential range of hyperactive AID variants with tunable activity, we permuted mutations previously shown to enhance AID activity (Supplementary Figure S1A). One cluster of mutations [C1] (AID-C1, Figure 1B), developed during a comprehensive structure-function analysis of AID, involved multiple changes in the ssDNA-binding loop, with R119G as a dominant activating mutation (22). A second cluster [C2] (AID-C2, Figure 1B), uncovered using an iterative selection strategy, included T82I as a dominant mutation (21). Prior work has also shown that a naturally occurring variant lacking the last exon (denoted as AID*, Figure 1B) also shows enhanced activity over wild-type AID (AID-WT) (29).

We generated a total of eight AID variants with candidate hyperactivating mutations. To measure deaminase activity, we employed the well-established rifampin mutagenesis assay (22, 30), whereby variants are overexpressed in *E. coli* and the frequency of acquired resistance to rifampin (Rif^R^) provides a reliable surrogate of deaminase activity (Figure 1C, Supplementary Table S1). In this assay, AID-WT increases Rif^R^ 11-fold relative to a catalytically-inactive AID-E58A, while AID* shows a further 4-fold increase over AID-WT. Adding single activating mutations (R119G or T82I) to AID* leads to a further increase, while we observed a 40-fold and 75-fold increase in Rif^R^ for AID-C1* and AID-C2*, respectively, relative to AID-WT. Strikingly, combining all mutations in AID-C12* results in a 127-fold increase in Rif^R^ (Figure 1C). The synergy between these independently derived hyperactivating mutations further supports the claim that AID has been naturally attenuated, likely to modulate risks posed to the genome, and that AID can be further engineered to improve its catalytic activity.

The development of a spectrum of AID variants positioned us to systematically explore the relationship between intrinsic deaminase activity and base editor outcomes. To this end, we incorporated the AID variants into BE3 constructs, creating a range of potentially hyperactive AID base editors (hBEs). To measure editing activity, we employed a reporter cell line with a single copy of destabilized GFP (*d2gfp*) integrated into HEK293T (26). Editing can be tracked by flow cytometry using an sgRNA that targets *d2gfp* to either generate indels with WT Cas9 or a nonsense Q158* mutation with base editing (Figure 1D, Supplementary Table S1). Using this assay, WT Cas9 editing is efficient, with 85±4% of GFP+ reporter cells converted to GFP^off^. With the same sgRNA, minimal GFP inactivation was observed with AID-WT-BE3. In line with our hypothesis, however, GFP inactivation significantly increases with hBEs (Figure 1D, Supplementary Figure S1B). Editing by AID*-BE3 results in 22±7% GFP^off^, with further enhancement by either R119G or T82I mutants. AID-C1*-BE3 and AID-C2*-BE3 showed 26±4.8% and 35±8.6% GFP^off^, respectively, while the full suite of synergistic hyperactivating mutations in AID-C12*-BE3 yields 53±2% GFP^off^. To integrate across the series, we plotted Rif^R^ frequency, a measure of intrinsic deaminase activity, against GFP^off^ frequency, a measure of BE activity (Figure 1E). An overall strong and positive correlation (r = 0.89) suggests that the intrinsic deaminase activity can be rate-limiting in the complex multi-step BE reaction.

### Deaminase activity shapes editing window

To better understand the mutational footprints of hBEs, we next deep-sequenced the *d2gfp* locus for five constructs spanning a range of intrinsic deaminase activities. With hBE constructs, target cytosine conversion (C > T/G/A/indels) increased relative to AID-WT-BE3 but appeared comparable across constructs (Supplementary Figure S1C, Supplementary Table S2), indicating that increased *d2GFP* inactivation did not exclusively come from nonsense mutation at the Q158 codon. When broadening analysis, we observed a striking pattern with multiple mutations across the locus, including C>T edits on both DNA strands (Figure 2A – Left, Supplementary Table S3). To help organize observations, we define NTS editing as C>T editing on the top strand (schematic in Figure 1A), and TS editing as C>T editing on the bottom strand, which is read as G>A mutations on the top strand, with no other mutations occurring above background levels. With hBEs, we observed both NTS (C>T) and TS (G>A) editing events, including outside the traditional BE window. At the level of individual DNA strands mutations also appear clustered tracking with increasing deaminase activity.

**Figure 2.**
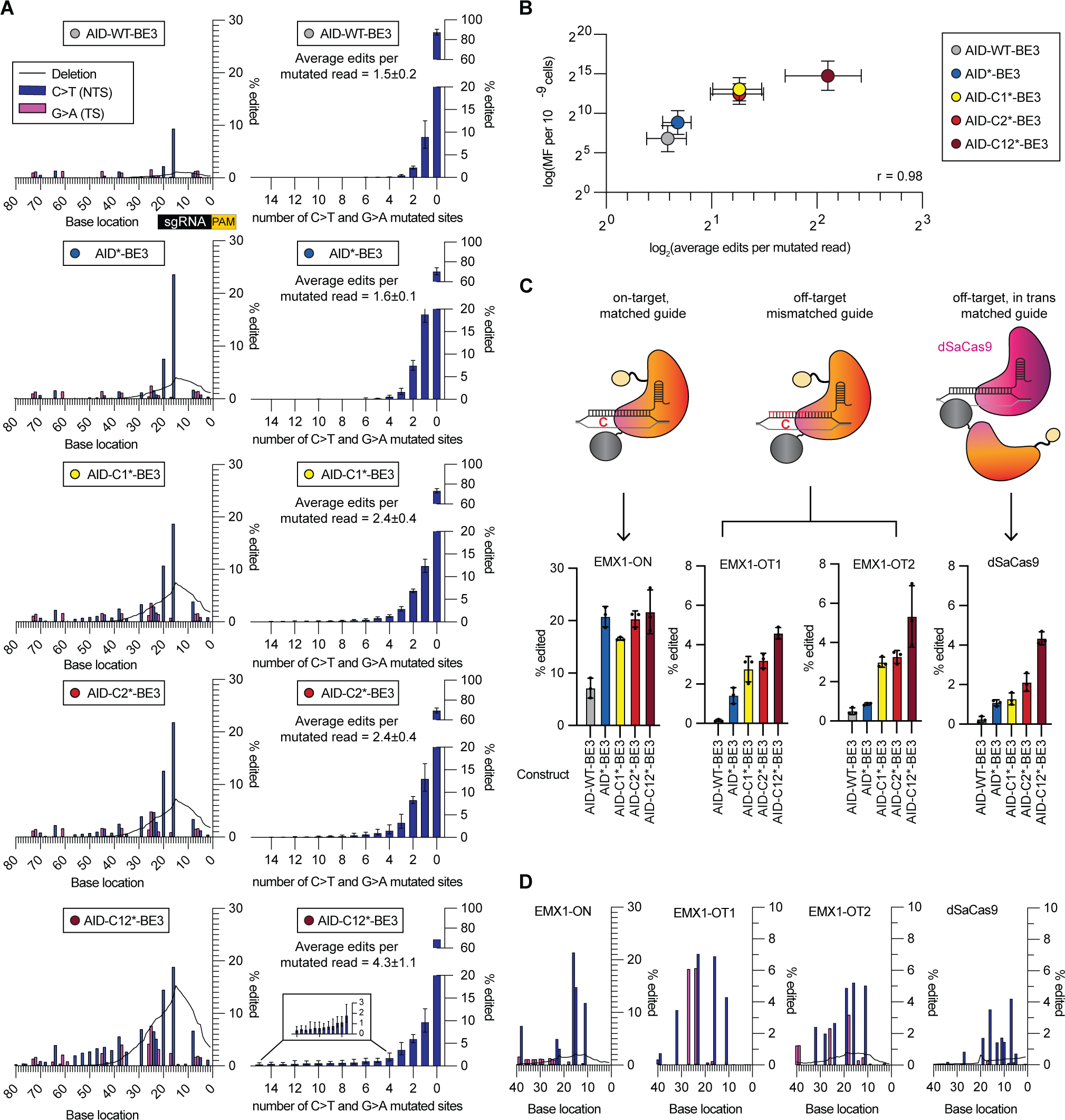
Hyperactive AID and Cas9 cooperativity establish hBE mutational footprints. (A) Left – Editing footprints of hBEs across the *d2gfp* locus for each condition. The sgRNA protospacer is located at bases 0-20, with the target C of the Q158 codon at position 16. The PAM is not shown (bases −1 to −3). Data are averaged across three replicates with individual values provided in Table S3. Right – Number of C>T and G>A mutations per mutated read for each condition (mean and standard deviation, n=3) with individual values provided in Table S4. (B) Correlation plot of intrinsic DNA deaminase activity as measured by Rif^R^ vs. average number of C>T and G>A mutations per mutated read. (C) Shown are deep sequencing C>T conversion efficiency of the target cytosine for three different Cas engagement modes: at the on-target site (EMX1-ON), two sgRNA-dependent off-target sites (EMX1-OT1 and EMX1-OT2) and one sgRNA-independent off-target site (dSaCas9). (D) Editing footprints for AID-C12*-BE3 across loci for data in (C) with individual values provided in Table S3.

To dissect the observed trends, we assessed the mutational load of the *d2GFP* locus after hBE treatment by quantifying the average number of C>T and G>A edits per mutated read (Figure 2A – Right, Supplementary Table S4). Cells treated with AID-WT-BE3 and AID*-BE3 harbored similar mutational loads, 1.5±0.2 and 1.6±0.1 mutations per read (mutations/read) respectively; however, AID*-BE3 reported higher activity on all sites in the *d2gfp* locus. AID-C1*-BE3 and AID-C2*-BE3, which have comparable intrinsic deaminase rates, both generate 2.4±0.4 mutations/read, while AID-C12*-BE3 was able to generate on average 4.3±1.1 mutations/read. Notably, we observed reads with as many as 15 combined C>T and G>A mutations/read with AID-C12*-BE3, a feature suggestive of its capacity to be a powerful diversifying base editor. When the average mutations/read was plotted against Rif^R^ rates, there was a strong correlation (r = 0.98) between intrinsic DNA deaminase activity and the overall mutational load (Figure 2B). To explore the generality of this result, we extended our analysis to a second genomic locus, *TDG,* and replicated our key findings, with tunable mutation loads and a broad editing footprint as a function of intrinsic deaminase activity (Supplementary Figure S2, Supplementary Tables S2 and S3). While we observed that increased efficiency broadens the editing window, our result provides a strong corollary to prior studies where stunting deaminase efficiency narrowed the editing window and increased precision (13). Our results also highlight that increasing AID catalytic activity not only increases BE activity, but also elicits the native processivity of AID, simulating its behavior on the Ig locus at sites of our preference.

### Cas9 engagement impacts DNA deaminase strand capture

There are multiple hypotheses addressing how AID targets both Ig strands in B-cells (31, 32) including antisense transcription (33, 34), DNA topology (35), and R-loop collapse by endogenous RNAse H digestion (36), all of which would expose both strands as ssDNA. However, how Cas9 binding can potentially create the right environment for AID to access either the NTS or TS has not yet been studied. Given clear signatures of both NTS (C>T) and TS (G>A) editing in our constructs, we next aimed to explore the DNA deaminase ‘strand capture’ step by perturbing Cas9 engagement. We designed our experiment around a method known as the R-loop assay, which allowed us to probe DNA deaminase activity at three sites with varying Cas9 engagement modes simultaneously (37). In this assay, HEK293T cells are co-transfected with three plasmids encoding the hBE construct, an *EMX1*-targeting sgRNA, and a catalytically-dead *S. aureus* Cas9 (dSaCas9) together with its sgRNA targeting a totally unrelated genomic locus. Using this system, the impact of Cas9 engagement on editing can be studied at (1) the on-target site (*EMX1*) with matched sgRNA, (2) known off-target sites (*EMX1*-OT1/2) where sgRNA is mismatched, as it is not fully complementary to the off-target sequences, and (3) the dSaCas9 target site where an R-loop is generated *in trans* and independent of the SpCas9-based BE (Figure 2C – Top).

As anticipated based on *d2GFP* and *TDG* results, while editing at the target C of the on-target *EMX1* locus plateaued (Figure 2C - Bottom), we observed a consistent increase in the average total C>T and G>A mutations per read across hBE constructs (Supplementary Figure S3). By contrast, at *EMX1*-OT1/2, AID-WT-BE3 showed near background levels of editing while hBE constructs show a consistent increase as a function of deaminase hyperactivation (Figure 2C - Bottom, Supplementary Figure S4). The altered pattern was even more striking when analyzing NTS (C>T) versus TS (G>A) mutations. At sgRNA-dependent off-target sites, unlike at the on-target site, TS (G>A) mutations were equally as prevalent as NTS (C>T) mutations for AID-C12*-BE3 (Figure 2D, Supplementary Figure S4), raising the possibility that sgRNA mismatches could permit AID capture and deamination of the TS. The R-loop generated *in trans* by dSaCas9 provided yet another stark contrast. While increased intrinsic deaminase activity still correlated with NTS mutations, TS mutations were nearly undetectable. These results suggest a high degree of cooperativity within BE3, and a clear and novel finding regarding the impact of the Cas9-sgRNA complex on deaminase activity. When the two DNA binding domains, Cas9 and AID, are acting *in cis*, AID can access not only the NTS, but also the TS, and this activity increases significantly with sgRNA mismatches. However, when the DNA deaminase acts *in trans* at an R loop generated by dSaCas9 with a perfectly matched sgRNA, the TS appears inaccessible.

Intrigued by these results, we wondered if the differences observed between the *in cis* and *in trans* behavior could also be impacted by the fact the *in cis* site is nicked, while the off-target R-loop site is not. We therefore repeated matched R-loop assay experiments for AID-C12*-BE3 employing either dSaCas9 or a SaCas9 TS-nickase (SaCas9-D10A, nSaCas9). In this comparison, *in cis* editing by AID-C12*-BE3 at the on-target *EMX1* locus remained the same, as expected (Supplementary Figure S5, Supplementary Table S5). However, we did observe a small increase in NTS mutations with the R-loop generated by nSaCas9 relative to dSaCas9, likely attributable to the directed repair of the nicked strand when using nSaCas9. Nonetheless, with either dSaCas9 or nSaCas9 involved in R-loop formation, the TS still appeared inaccessible, supporting our observation that perfectly-matched sgRNAs minimize TS editing.

Given that off-target editing increased with hBEs, we also considered whether the higher deamination activity could be toxic to mammalian cells. To assess this possibility, we evaluated both total cell viability and the induction of DNA damage responses after hBE editing. Induction of the DNA damage response was measured by phosphorylated γH2AX as a marker of dsDNA breaks, and either the DNA deaminase APOBEC3A or hydroxyurea treatment were used as positive controls for comparison (38, 39). Our results indicate no significant increase in cytotoxicity with any of the BE constructs (Supplementary Figure S6, Supplementary Table S6). When comparing the inactive AID(E58A)-BE3 to hyperactive AID-C12*-BE3, there was, however, a modest but measurable increase in the DNA damage response marker. This increase was dependent upon the presence of a targeting sgRNA, suggesting that the combination of nCas9 with hyperactive AID can increase in dsDNA breaks, a result that aligns with our sequencing results where higher levels of indels were observed with the hBEs.

### Cas9 nuclease activity shapes the landscape of NTS vs. TS mutations

Intrigued that hBEs have broadened editing windows and generate TS mutations, we first aimed to increase hBE activity by moving AID-C12* from the BE3 into the BE4max scaffold (10). As predicted with this change, when targeting the *d2GFP* locus, AID-C12*-BE4max led to a 1.8-fold increase in target base C>T conversion and a 2.6-fold decrease in deletions when compared to AID-C12*-BE3 (Supplementary Figure S7, Supplementary Table S7). As a second possible modification to the BE scaffold that could impact activity, we also assessed if the editing window could be further broadened by using a longer linker between AID-C12* and nCas9. To explore this possibility, we replaced the original BE4max linker (32 amino acid long SGGS-XTEN-SGGS linker) with an 80 amino acid long XTEN80 linker that has been employed in some epigenome editors (24). In comparing the standard and extended linkers in the BE4 scaffold, we observed that the longer linker retained high activity inside the protospacer but led to a modest decrease in editing activity outside of the protospacer for both NTS and TS (Supplementary Figure S8, Supplementary Table S8). These results suggest that the ‘capture’ of the region outside of the traditional window by AID is likely to be influenced by linker length in a complex manner, such that proximity is important for optimal engagement. Given the higher levels of editing with the traditional linker from the BE4max construct, we focused further analysis on this construct.

Given the overall improved performance when targeting the *d2GFP locus*, we next characterized AID-C12*-BE4max across a broader array of genomic sites. We targeted seven independent genomic loci (Supplementary Table S9), allowing us to derive an aggregate profile of C base editing on either strand across an 80 bp region flanking the PAM (Figure 3A, Supplementary Table S10). Similar to traditional BEs, in these aggregate profiles, maximal NTS (C>T) editing by hBEs occurs inside the protospacer located 10-20 bp away from the PAM, with a maximum of ∼25-38% editing reached across the seven loci (Figure 3A). Upstream of the maximal editing window, editing tapers away but can be detected up to 80 bp away from the PAM sequence. However, downstream of the maximal editing window, editing quickly diminishes with mutations appearing undetectable further downstream of the PAM. This observation reveals that Cas9-binding to the PAM obstructs bidirectional AID processivity, skewing mutations to the region upstream of the PAM only.

**Figure 3.**
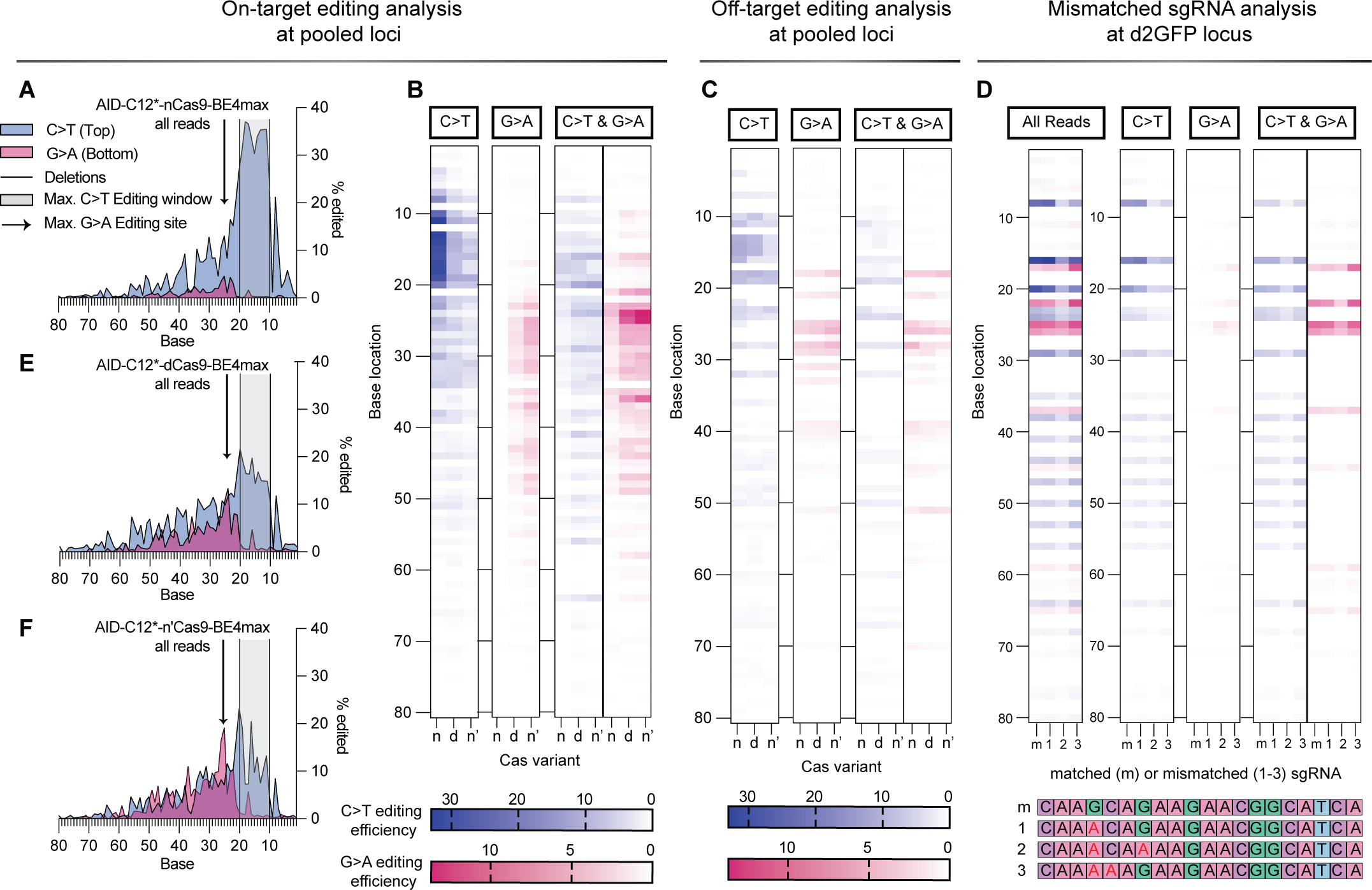
Perturbing Cas9 cleavage activity skews BE outcomes. Shown are the averaged editing footprints and heat maps from strand specific analysis of on-target or off-target sites. For AID-C12*-BE4max, the editing frequency is averaged across seven loci in (A), presented with the sgRNA protospacer located at bases 0-20, while PAM is not shown (−1 to −3). The maximal C>T and G>A editing windows are noted with shading and an arrow, respectively. The heat maps of strand-specific analysis showing C>T only, G>A only and simultaneous C>T/G>A mutations are provided in (B) in the corresponding left columns (n, nCas9). Similar heat maps averaged across four off-target sites is provided in (C) in the left columns (n, nCas9). Data for individual on-target and off-target sites are given in Table S10 and Table S11, respectively. (D) Matched (m) and mismatched sgRNAs (1–3) are shown at bottom, with mismatches in red text. The heatmaps for NTS and TS editing at the d2GFP locus with AID-C12*-BE4max are shown for total edits and segregated by reads with C>T only, G>A only, and simultaneous C>T/G>A mutations. Data for individual replicates are provided in Table S12. (E) and (F) Analogous to the results in (A), the on-target averaged editing footprints across seven loci for AID-C12*-dCas9-BE4max and AID-C12*-n’Cas9-BE4max are shown. The heatmaps are similarly shown in (B) and (C), with the dCas9 construct results shown in the middle columns and the n’Cas9 (n’) construct results at right.

For G>A transitions, which occur upon AID engagement with the TS, we observe a different pattern. Maximal editing occurs outside the protospacer, peaking ∼21-28 bp away from the PAM and reaching ∼15% activity on average across the seven loci (Figure 3A). This finding suggests that sgRNA-engagement effectively blocks AID access to the TS in the protospacer region. G>A edits also taper away, but can be detected at >1% as far as 50 bp upstream from the PAM.

To better understand the mechanistic basis for TS mutations, we next analyzed the pattern of mutations in individual sequencing reads (Figure 3B, left most column ‘n’ for nCas9). Interestingly, the majority of C>T transitions occurred alone. In contrast, G>A transitions were almost exclusively observed in reads that also harbored C>T transitions, and not in isolation. We hypothesized that NTS (C>T) mutations could happen preferentially during a BE ‘first encounter’. After DNA replication leads to the fixture of these C>T mutations in the NTS, a ‘second encounter’ with a BE can occur. The presence of mismatches in the sgRNA-binding site arising from the first encounter could then increase access to the TS for editing, which would result in both C>T and G>A transitions on the same read. To further substantiate this model, we analyzed the sgRNA-dependent off-target effects at both *EMX1* and *FANCF* off-target sites (Supplementary Figure S9 – Left, Supplementary Table S11). These sites are marked by multiple PAM-distal sgRNA mismatches that would be present in the first encounter. In line with our hypothesis, we observe an increase in G>A mutations overall. More specifically, however, strand-specific analysis revealed a marked increase in G>A only mutations, indicating that in the presence of mismatches, TS editing alone is increased and can happen in a first encounter with AID-C12*-BE4max (Figure 3C, left most column ‘n’ for nCas9). As an orthogonal method of validating that TS editing is increased in the presence of mismatches, we designed three sgRNAs (1–3) to mimic mismatches that would commonly arise between the genomic DNA locus and the *d2GFP* targeting sgRNA at the second encounter (Figure 3D, Supplementary Table S12). Unlike the matched sgRNA (m), using these mismatch-containing sgRNAs more readily yielded G>A mutations in the absence of C>T mutations, albeit at low levels. Interestingly, the greatest yield of G>A only mutations was observed with mismatched sgRNA-3, which contains two consecutive mismatches at the target cytosine, relative to the single mismatch in sgRNA-1 and the two non-consecutive mismatches in sgRNA-2. These observations further support our hypothesis that TS editing can occur at on-target sites in the presence of a mismatch that is encoded in either the sgRNA or in the target locus.

As our results suggested two distinctive aspects of the hBEs, activity outside the traditional protospacer editing window and editing of the TS, we considered whether intrinsic features of AID beyond the observed processivity could be detected in the editing footprint. In vitro and in vivo experiments have shown that AID deamination efficiency is strongly influenced by upstream bases, with a preference for WR<u>C</u> (W = A/T; R = A/G) motifs as hotspots for mutations (23, 40-42). To explore if sequence preferences manifest in the hBE editing window, we reanalyzed the mutational footprint by classifying the target C bases on the TS and NTS strands as being hotspots (WRC), coldspots (SYC; S = C/G; Y = T/C) or neither (Supplementary Figure S10 – Left, Supplementary Table S13). We observe that there seems to be no preference for WRC motifs inside the protospacer, presumably because AID has continuous access to the R-loop generated by nCas9. At increasing distance from the protospacer, however, there is an increase in the editing of WRC hotspots when compared to SYC cold spots or other sequence contexts (average NTS C>T editing of 3.2% versus 0.6% for C bases 41-80 bp from PAM). These results align with our model, where the intrinsic preferences of AID play a larger role in editing regions where ssDNA generation is not enforced by nCas9 binding.

### Manipulating Cas9 cleavage activity skews BE outcomes and yields a proficient diversifying editor

In BE4max, nCas9 nicking of the TS increases the probability of NTS mutations due to the preferential recognition and excision of the mismatched base present on the nicked DNA strand. As such, nick-associated repair could obscure the extent of NTS versus TS editing. To further explore how Cas9 could cooperatively alter base editing outcomes, we next exchanged nCas9 in the AID-C12*-BE4max construct with either dCas9 or Cas9-H840A (n’Cas9), a variant that nicks the NTS rather than the TS, which could increase the probability of TS mutations. To construct an overall targeting profile, we employed these variants to edit the same seven genomic loci as with AID-C12*-BE4max.

As anticipated when losing TS nicking with the dCas9 construct, we observe a decrease in overall C>T editing activity, with a range of ∼11-20% maximal editing across the seven loci (Figure 3E). By contrast, we observed an increase in NTS (C>T) transitions, which localizes to the same window (∼21-28 bp upstream of PAM) as observed with the nCas9 construct. Interestingly, however, strand-specific analysis, demonstrated that G>A edits were now readily detectable by themselves in the absence of C>T edits (Figure 3B, middle column ‘d’ for dCas9). These results support a model where capture and mutation of the TS can occur in normal BE4max constructs in the first encounter, but such mutations can be reversed by the repair initiated at the nick and are unmasked with the use of dCas9.

Based on the established model, we anticipated that AID-C12*-n’Cas9-BE4max might be a particularly powerful diversifying base editor. Analyzing across the seven target loci, we find that G>A edits further increase relative to the dCas9 construct, reaching levels that rival those of C>T editing (Figure 3F). Strand-specific analysis revealed a nearly equal prevalence of sequences with C>T alone, G>A alone or dual C>T and G>A edits, with detectable mutations at a frequency of >1% extending over a 65 bp window (Figure 3B, right most column ‘n’’ for n’Cas9). Thus, AID-C12*-n’Cas9-BE4max appears to offer the broadest diversifying window to date for a DNA deaminase-linked BE construct. It is worth noting that we observe the same trend with both dCas9 and n’Cas9 hBEs as with nCas9, where there is no sequence context preference inside the protospacer, but WRC hotspots are preferred outside of the traditional base editing window (Supplementary Figure S10 – middle and right, Supplementary Table S13).

Returning to our model, we also examined editing at the *EMX1* and *FANCF* off-target sites. Consistent with the model, the dCas9 and n’Cas9 editors show a pattern where G>A only transitions are relatively increased due to the sgRNA mismatches further facilitating TS editing in the first encounter (Figure 3C, middle and right columns, Supplementary Figure S9 – middle and right, Supplementary Table S11). Hence, our results demonstrate that modulating Cas9 catalytic activity in concert with hyperactivation of AID can mimic somatic hypermutation across highly diverse loci with processive, targeted diversification across strands.

## DISCUSSION

In this study, we substantiate a complete mechanistic model for the BE mechanism that accounts for Cas9 engagement, strand capture by AID, deamination, and DNA repair (Figure 4). Specifically, our study reveals several previously unappreciated means whereby Cas9 and AID collaborate to promote BE outcomes.

**Figure 4.**
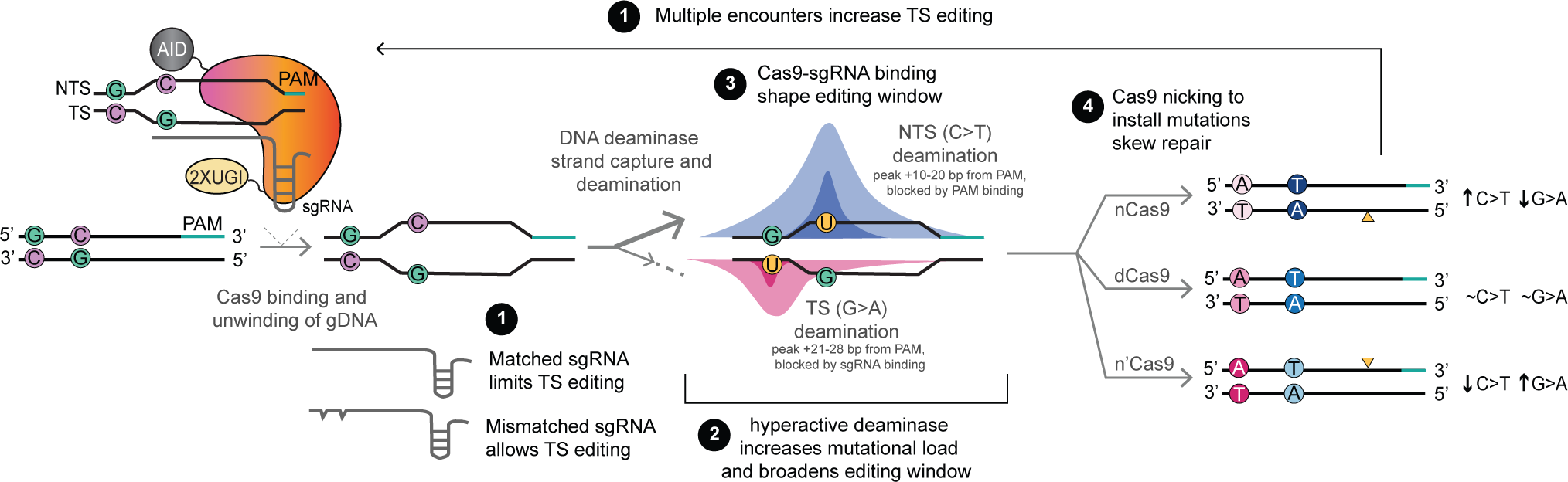
Cooperativity between Cas9 and AID shapes BE outcomes. In the mechanistic model, Cas9 and the DNA deaminase work together to determine BE outcomes in multiple manners. 1-During an initial hBE encounter, the nature of the R-loop determines whether AID binds the NTS or TS. Matched sgRNA binding promotes NTS editing, while mismatches, which occur at either off-target sites or during second encounters at on-target sites, promote increased TS editing. 2-Once engaged with a strand, AID can act processively to introduce multiple C>U mutations, with the number of edits and the size of the editing window depending on the intrinsic deaminase activity. 3-Cas9 binding defines the borders of editing, with NTS edits peaking inside the protospacer and TS edits peaking outside the protospacer. 4-Finally, the presence of a nick on one strand can increase editing at the other strand. At each step in the mechanism, activity is thus determined by the cooperative actions of the two DNA modifying proteins, Cas9 and AID, acting *in cis*.

First, we show that strand capture by the DNA deaminase is impacted by the nature of the R-loop generated by Cas9. At on-target sites, binding of a matched sgRNA to the TS results in preferential exposure of the NTS, leading to its capture by the deaminase and subsequent generation of C>T mutations. By contrast, at off-target sites, in the presence of sgRNA-TS mismatches, the TS can more readily be accessed and deaminated, resulting in an increase of G>A mutations (reading from the perspective of the NTS). The lack of G>A only mutations detected with AID-C12*-BE4max during a first BE encounter at on-target sites provides orthogonal evidence to support this claim. After the initial introduction of NTS (C>T) mutations, sgRNA mismatches are created, increasing TS capture in a second encounter, and leading to detectable G>A mutations in the presence of the already established C>T mutations. The introduction of sgRNA mismatches at a target site or the presence of such mismatches at off-target sites can allow for detectable G>A mutations in the absence of C>T mutations. When taken in the context of studies showing that Cas9 accommodates PAM distal mismatches (43), our data suggest that the R-loop formed in such cases is dynamic, allowing for capture of the sgRNA-target strand by the DNA deaminase.

Second, we show that once the strand has been captured, the overall base editing activity and the editing window can be tuned as a function of the intrinsic activity of the DNA deaminase. By using the same AID scaffold and variants spanning several orders of magnitude in Rif^R^ activity, we show that intrinsic deaminase activity can govern the overall BE reaction. As activity is further increased, the intrinsic properties of DNA deaminases, specifically AID’s processivity, are evident with strand-specific clustered mutations that can extend far outside of the canonical editing window opened by Cas9. Additionally, sequence preferences intrinsic to the deaminase can be observed to take on more importance at increasing distance from the protospacer. These observations align with prior work suggesting that decreasing the editing activity of the DNA deaminases can increase the precision of editing (9, 13), as well as studies that show that bystander edits arise with slower kinetics than on-target editing (44).

Third, as with strand capture, we show that there is cooperativity between Cas9 and AID in establishing the mutational footprint. On the NTS, the PAM binding site serves as a barrier preventing downstream mutagenesis. By contrast on the TS, matched sgRNA binding largely blocks mutations on the protospacer, leading to distinctive maximal peaks for C>T versus G>A mutations. In the context of structural studies capturing intermediates in R-loop formation (45), our work highlights the strength of the PAM interaction as a stable barrier obstructing AID processivity on the non-target strand, with sgRNA binding as a less stringent, but significant barrier to target strand editing.

Fourth, we demonstrate that in concert with the DNA deaminase, Cas9’s catalytic activity further contributes to editing outcomes. Prior work has shown that moving from dCas9 in BE1 to nCas9 in BE3 leads to higher C>T editing activity (2). Here, by changing nCas9 to dCas9 in AID-C12*-BE4max we also show that nicking of the TS results in the suppression of rare TS (G>A) editing events. Changing to n’Cas9 results in an even higher degree of TS mutations alone, although the fact that the nick is positioned downstream of the mutations likely limits the ability of DNA replication to install mutations as readily. These results highlight the importance of downstream DNA repair pathways in either resolving or embedding mutations, aligning with recent work that highlights the cell cycle dependence of base editing outcomes (46).

We conclude by highlighting how dissecting the molecular mechanisms of individual steps involved in a complex multi-step BE reaction offers an effective means to develop new tools. In our case, by hyperactivating AID, altering Cas9 activity via nickase manipulation and studying different modes of Cas9 engagement at off-target sites, we generated a novel diversifying base editor (AID-C12*-n’Cas9-BE4max) that produces C>T only, G>A only, and concurrent C>T and G>A mutations with nearly equal efficiency over a window of more than 65 bp. These features rival alternative approaches that have been taken to achieve diversification, such as dual ABE/CBE strategies, recruitment of multiple deaminases to the same site, or targeted action of error-prone polymerases (47–50) Taken together, our study highlights how mechanistic inquiries can continue to drive the discovery and expansion of the genome editing toolbox.

## DATA AVAILABILITY

Novel plasmids generated in this study will be deposited to Addgene. Next-Generation Sequencing (NGS) data raw fastQ files were deposited to the National Center for Biotechnology Information (NCBI) - BioProject: PRJNA891058. Access for the reviewers can be obtained via: https://dataview.ncbi.nlm.nih.gov/object/PRJNA891058?reviewer=a7kcrcj6kotiqlacphksa5puh2.

Unprocessed NGS data from CRISPResso2 is available in the Supplementary Data. Author’s original code and the NGS data from CRISPResso2 processed with the code is available in the Supplementary Data.

Further information and requests for reagents should be directed to and will be fulfilled by Rahul M. Kohli and Junwei Shi.

## CODE AVAILABILITY

Original Codes to analyze NGS data in Rstudio and their descriptions are supplied in the Supplementary Data.

## SUPPLEMENTARY DATA

Supplementary Data are available at NAR online.

## AUTHOR CONTRIBUTIONS

K.N.B., J.S., and R.M.K. conceived of the approach. K.N.B., A.B., J.S., and R.M.K. designed the research. K.N.B., A.B., J.C.S, P.F.B., J.P. and K.S.G. performed experiments and analyzed data. K.N.B., A.B., J.G., P.F.B. and N.H.E. performed computational and statistical analysis. The manuscript was drafted by K.N.B., revised by J.S., and R.M.K., with added input and approval from all authors.

