## Supplementary Tables for "Cooperativity between Cas9 and hyperactive AID establishes broad and diversifying mutational footprints in base editors"

*Supplementary Data Figures*

Supplementary Figure S1: AID variants and hBEs flow cytometry plots ............................S2
Supplementary Figure S2: hBE activity at the *TDG* locus ..................................................S3
Supplementary Figure S3: hBE activity at the *EMX1* locus ................................................S4
Supplementary Figure S4: Off-target hBE activity .............................................................S5
Supplementary Figure S5: Modified R-loop assay with nSaCas9........................................S6
Supplementary Figure S6: Cytotoxicity and DNA damage by hBEs..................................S6
Supplementary Figure S7: BE3 and BE4max comparison ..................................................S7
Supplementary Figure S8: BE4max standard linker versus extended linker comparison .....S7
Supplementary Figure S9: Off-target editing analysis at pooled loci...................................S8
Supplementary Figure S10: Hotspot analysis of hyperactive AID base editors...................S8

*Supplementary Tables Legends*.........................................................................................S9-S10

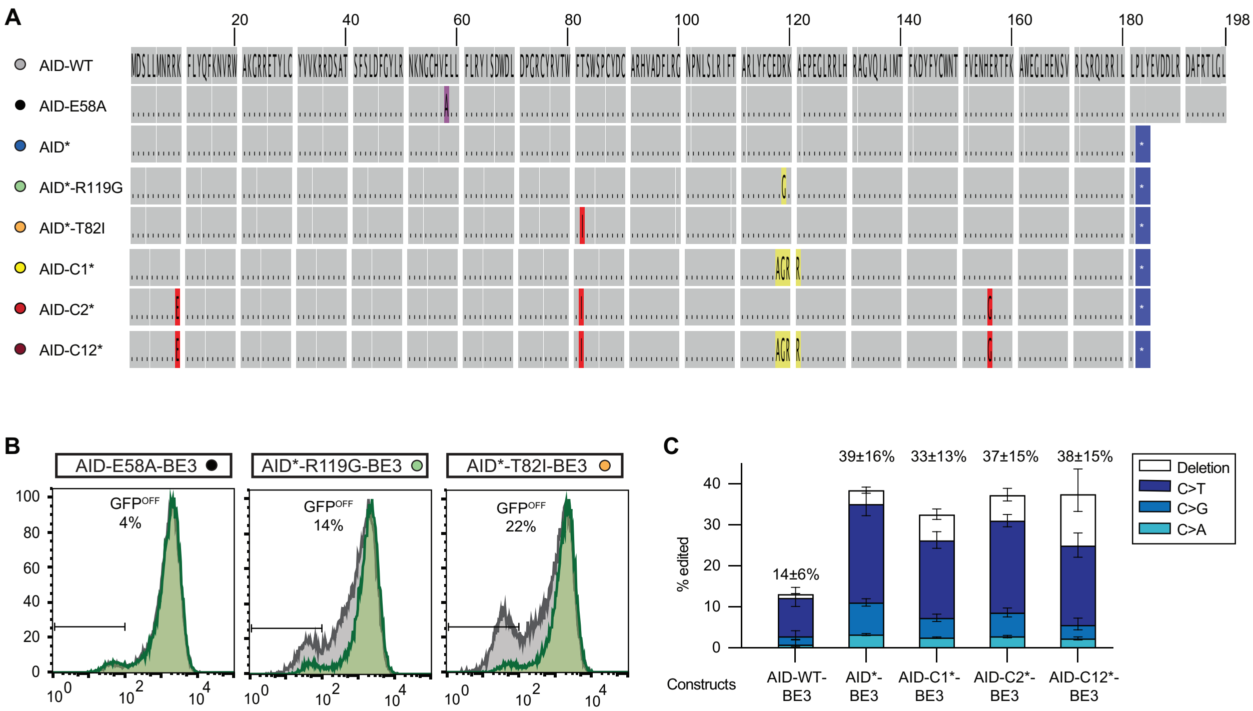


**Supplementary Figure S1: AID variants and hBEs flow cytometry plots. (A)** Schematic of AID protein sequence detailing the mutations of each variant deaminase tested. **(B)** Representative flow cytometry histograms associated with transfection of hBEs in HEKT293T-d2GFP cells for constructs not shown in Fig. 1. **(C)** Deep sequencing shows conversion efficiency at the target C to T, G, A and deletions (mean and standard deviation, n=3) with individual values provided in Tables S2 and S3.

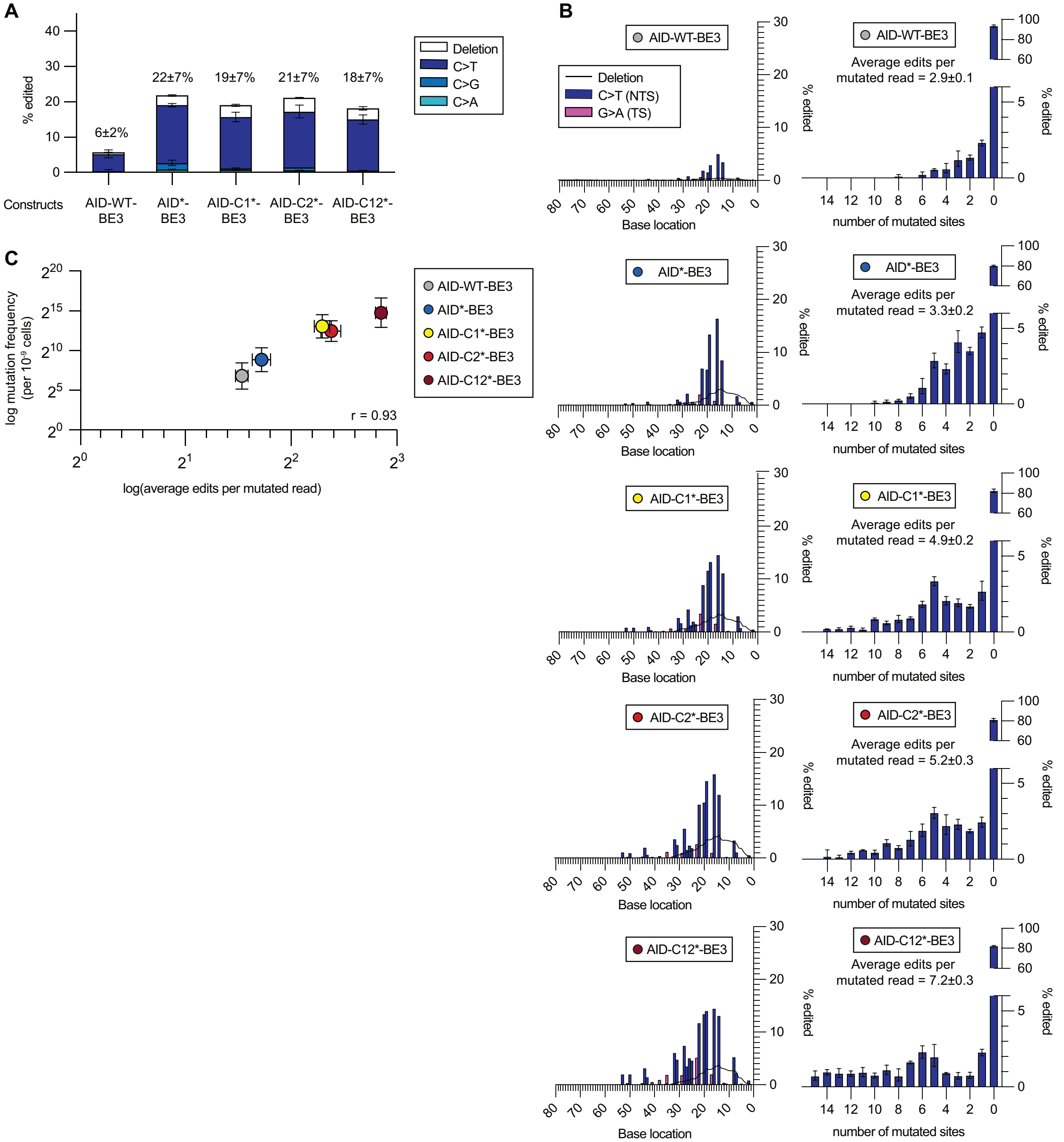


**Supplementary Figure S2: hBE activity at the *TDG* locus**. **(A)** Deep sequencing results with BE constructs demonstrating conversion efficiency at the target C to T, G, A and deletions (mean and standard deviation, n=3) with individual values provided in Table S2 **(B)** Left - Editing footprints of hBEs across the *TDG* locus for each condition. The sgRNA protospacer is located at bases 0-20. The PAM is not shown (bases -1 to -3). Data are averaged across three replicates. Right – Number of C>T and G>A mutations per mutated read for each condition (mean and standard deviation, n=3). Individual editing values provided in Tables S2 and S3. **(C)** Correlation plot of intrinsic DNA deaminase activity as measured by Rif$^R$ vs. average number of C>T and G>A mutations per mutated read.

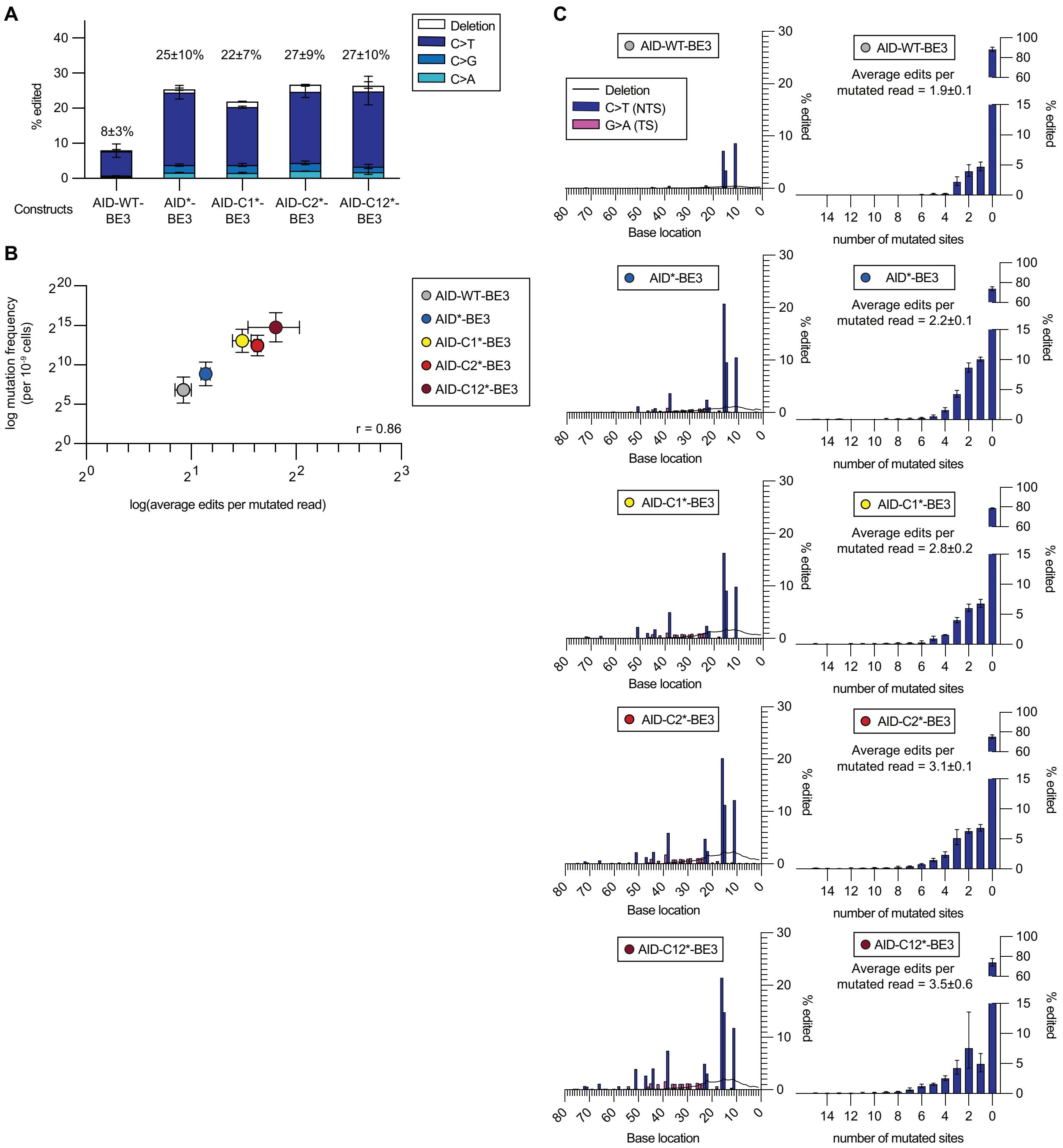


**Supplementary Figure S3: hBE activity at the *EMX1* locus. (A)** Deep sequencing results with BE constructs demonstrating conversion efficiency at the target C to T, G, A and deletions (mean and standard deviation, n=3) with individual values provided in Table S2. **(B) Left** - Editing footprints of hBEs across the *EMX1* locus for each condition. The sgRNA protospacer is located at bases 0-20. The PAM is not shown (bases -1 to -3). Data are averaged across three replicates. **Right** – Number of C>T and G>A mutations per mutated read for each condition (mean and standard deviation, n=3). Individual editing values provided in Tables S2 and S3. **(C)** Correlation plot of intrinsic DNA deaminase activity as measured by $Rif^R$ vs average number of C>T and G>A mutations per mutated read.

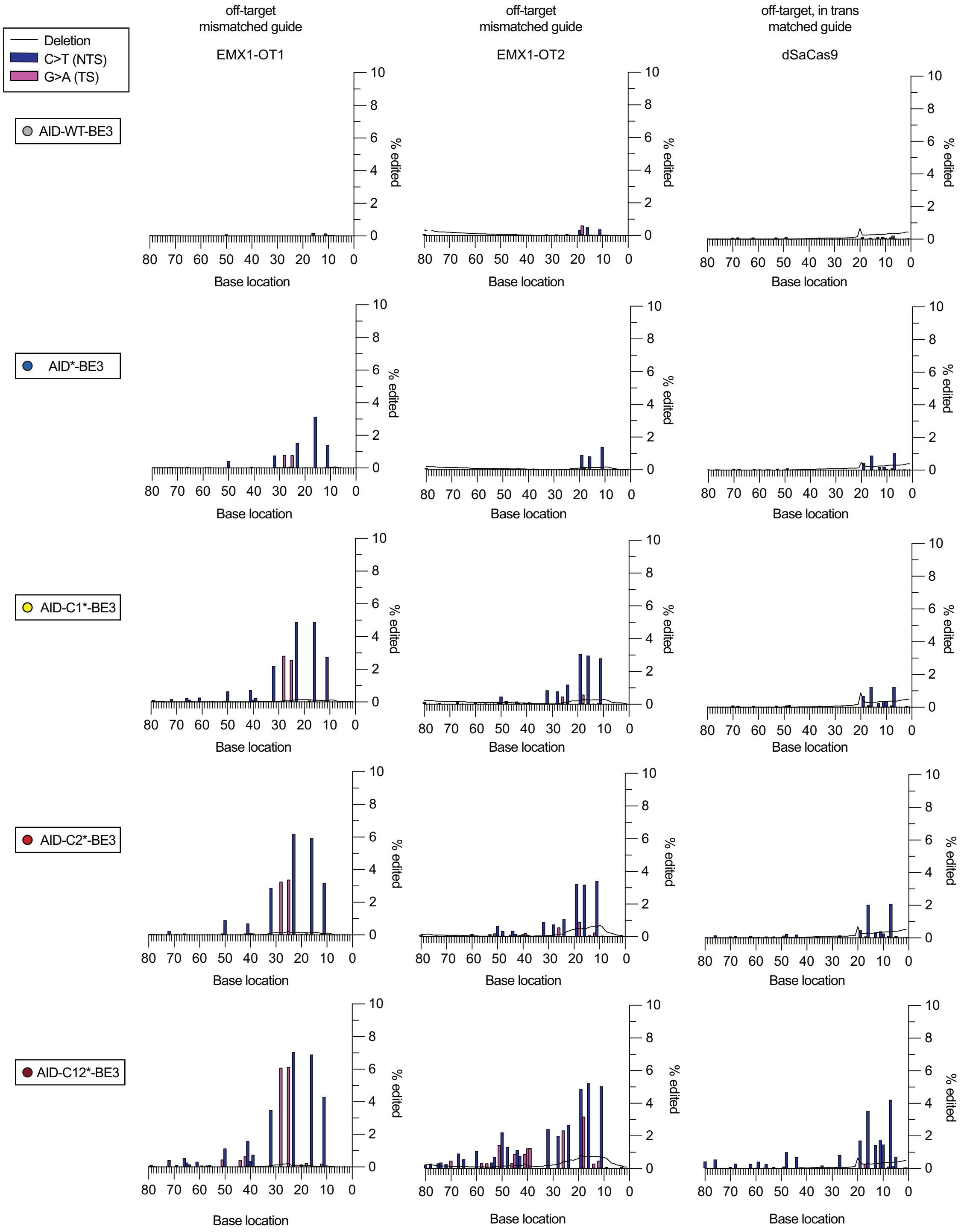


**Supplementary Figure S4: Off-target hBE activity.** Editing footprints of hBEs are shown across two sgRNA-dependent off-target sites (*EMX1-OT1* and *EMX-OT2)* and one sgRNA-independent off-target site (dSaCas9 binding site) generated by the R-loop assay. The sgRNA protospacer is located at bases 0-20. The PAM is not shown (bases -1 to -3). Data are averaged across three replicates. Individual editing values provided in Table S3.

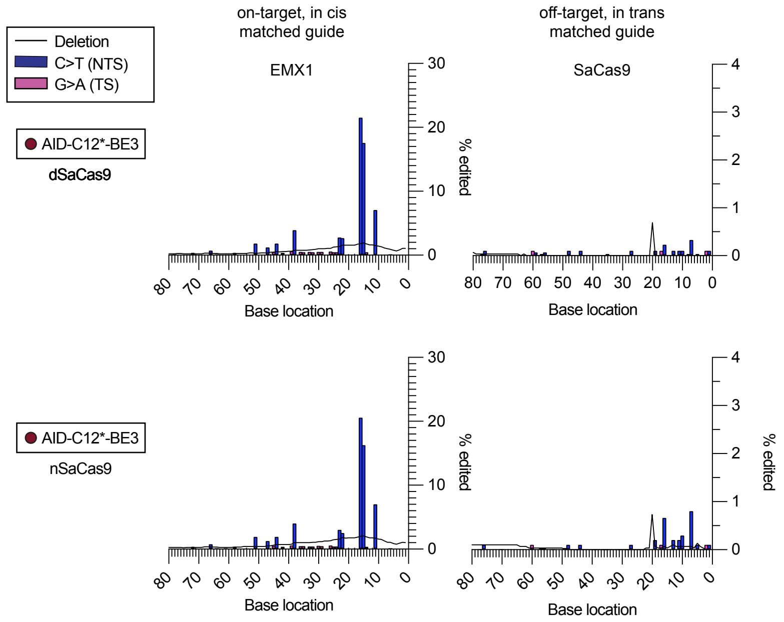


**Supplementary Figure S5: Modified R-loop assay with nSaCas9.** Editing footprints of AID-C12*-BE3 are shown at the EMX1 on-target site and an sgRNA-independent off-target site using dSaCas9 in the standard R-loop assay or nSaCas9 in the modified R-loop assay. The sgRNA protospacer is located at bases 0-20. The PAM is not shown (bases -1 to -3). Data are averaged across three technical replicates. Individual editing values provided in Table S5.

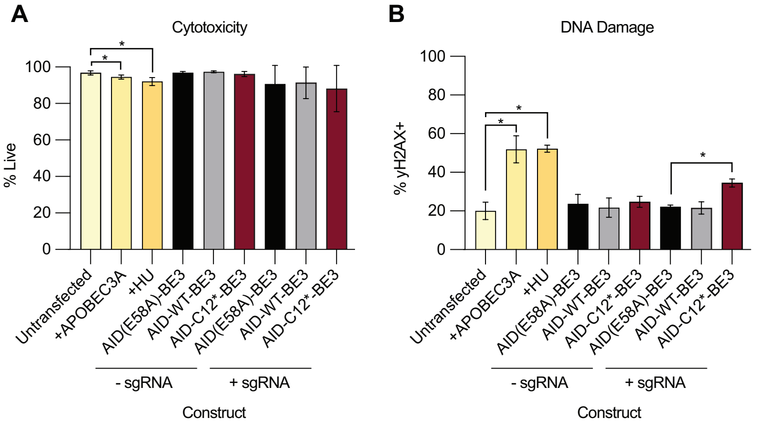


**Supplementary Figure S6: Cytotoxicity and DNA damage by hBEs. (A)** Analysis of cell viability and **(B)** DNA damage after base editing in the absence or presence of sgRNA. DNA damage is measured by quantification γH2AX. APOBEC3A, a highly active DNA deaminase, and hydroxyurea (HU), a potent inhibitor of DNA replication, were used as positive controls. A two-sided Mann–Whitney test was performed to compare between samples (*$P \leq 0.05$). Individual measurements and exact *P* values are provided in Table S6.

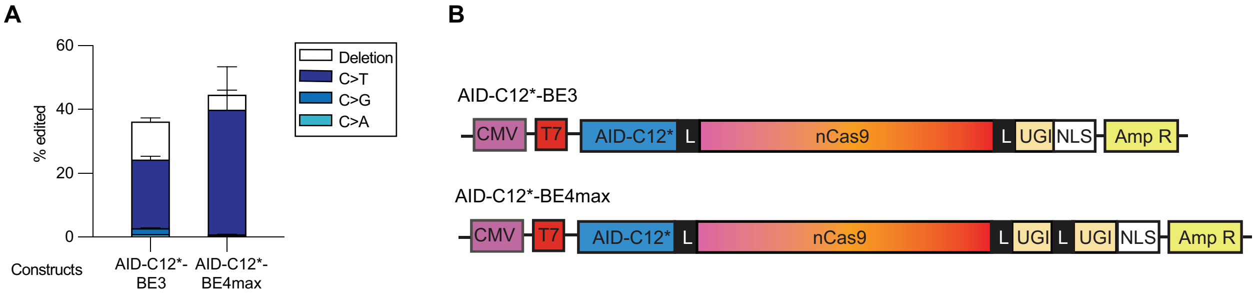


**Supplementary Figure S7: BE3 and BE4max comparison.** **(A)** Deep sequencing comparing target cytosine (C) conversion efficiency to T, G, A and deletions by BE3 and BE4max (mean and standard deviation, n=3) with individual values in Table S7. **(B)** Construct schematics for AID-C12*-BE3 and AID-C12*-BE4max.

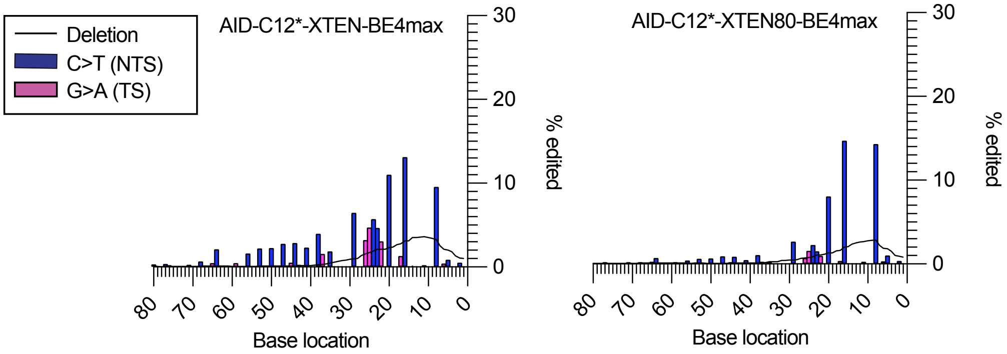


**Supplementary Figure S8: BE4max standard linker versus extended linker comparison.** Editing footprints of hBEs across the *d2GFP* locus for each condition. The sgRNA protospacer is located at bases 0-20. The PAM is not shown (bases -1 to -3). Data are averaged across three technical replicates. Individual editing values provided in Table S8.

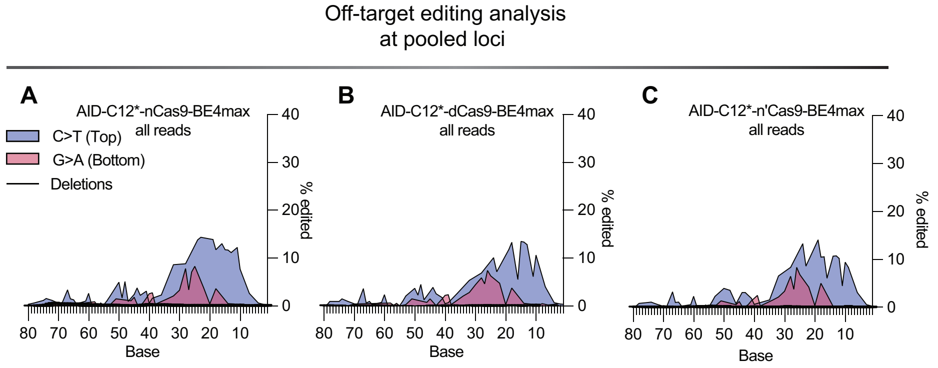


**Supplementary Figure S9: Off-target editing analysis at pooled loci.** Shown are the editing footprints from strand specific analysis averaged across four off-target sites with individual values provided in Table S11. The sgRNA protospacer is located at bases 0-20. The PAM is not shown (bases -1 to -3). The values for AID-C12*-BE4max are shown in (**A**), AID-C12*-dCas9-BE4max are shown in (**B**) and AID-C12*-n'Cas9-BE4max are shown in (**C**).

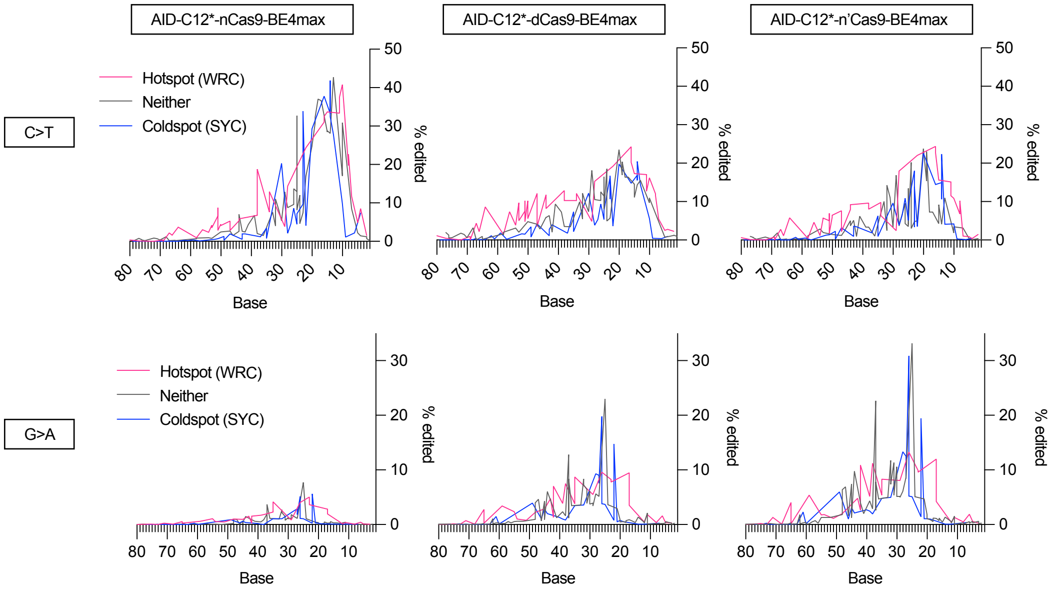


**Supplementary Figure S10: Hotspot analysis of hyperactive AID base editors.** Shown are the editing footprints from strand specific analysis averaged across seven loci for AID-C12*-nCas9-BE4max (left), AID-C12*-dCas9-BE4max (middle) and AID-C12*-n'Cas9-BE4max (right). Cytosine bases are classified as hotspots (WRC), coldspots (SYC) or neither. The sgRNA protospacer is located at bases 0-20. The PAM is not shown (bases -1 to -3). Complete data are provided in Table S13.

### SUPPLEMENTARY TABLES LEGENDS

**Supplementary Table S1: Rif mutagenesis and Flow cytometry assay data.** Data and statistical analysis for data shown in Fig. 1C, Fig. 1D, and Fig. 1E.

**Supplementary Table S2: Unprocessed d2GFP, TDG_g1, EMX1, target C.** Unprocessed CRISPResso2 base editing results from d2GFP, TDG_g1 and EMX1 sites, and statistical analysis on the target cytosine editing efficiency.

**Supplementary Table S3: Processed hBE editing.** Processed individual replicate data for how editing efficiency changes as the position of bases increases 5' away from the PAM at d2GFP, TDG_g1, EMX1, EMX1_OT1, EMX1_OT2 and dSaCas9 sites processed with original code "3AllTypes_Processivity".

**Supplementary Table S4: Mutational Load.** Overall mutagenic activity (both NTS and TS edits) of hyperactive AID base editors at d2GFP, TDG_g1 and EMX1 sites processed with original code "1Generalized_BarGraphMutatedSites"

**Supplementary Table S5: Unprocessed data from R-loop assay comparing dSaCas9 to nSaCas9.** Unprocessed CRISPResso2 base editing results from on-target EMX1 and off-target sites when assessing standard R-loop assay with dSaCas9 and modified R-loop assay with nSaCas9.

**Supplementary Table S6: Cytotoxicity and DNA damage after hBE expression.** Data and statistical analysis of cell viability (%Live) and DNA damage (γH2AX) for data shown in Supplementary Figure S6.

**Supplementary Table S7: Unprocessed BE3 vs BE4max, target C.** Unprocessed CRISPResso2 base editing results from the d2GFP site targeted with BE3 or BE4max, and statistical analysis on the target cytosine editing efficiency for data shown in Supplementary Figure S7.

**Supplementary Table S8: Unprocessed hBE editing of d2GFP with standard BE4max and BE4max with XTEN80 linker.** Unprocessed technical replicate data for base editing of the d2GFP locus with AIDC12*-XTEN-BE4max and AIDC12*-XTEN80-BE4max for data shown in Supplementary Figure S8.

**Supplementary Table S9: Sequencing Sheet.** Target gene, sgRNA sequence, target strand, sequencing primers, amplicon size, amplicon sequence and chromosomal location of all edited sites in this study.

**Supplementary Table S10: Processed Average Genomic Loci.** Processed average data for how editing efficiency changes as the position of bases increases 5' away from the PAM at seven genomic sites processed with original code "2Processivity_Average".

**Supplementary Table S11: Processed Average Off-Target Loci.** Processed average data for how editing efficiency changes as the position of bases increases 5' away from the PAM at four off-target sites processed with original code “2Processivity_Average”.

**Supplementary Table S12: Unprocessed and Processed hBE editing of d2GFP with mismatched gRNAs.** Unprocessed and processed individual replicate data for base editing of the d2GFP locus with AID-C12*-BE4max with matched (m) and mismatched sgRNA for data shown in Fig. 3D.

**Supplementary Table S13: Sequence context analysis of hyperactive AID base editors.** Reanalyzed mutational footprints for hBE constructs from seven averaged genomic loci where the target C bases on the TS and NTS strands were classified as being hotspots (WRC), cold spots (SYC) or neither for data shown on Supplementary Figure S10.
